# Transplanted Adult Neural Stem Cells Express Sonic Hedgehog *In Vivo* and Suppress White Matter Neuroinflammation After Experimental Traumatic Brain Injury

**DOI:** 10.1101/196170

**Authors:** Genevieve M. Sullivan, Regina C. Armstrong

**Affiliations:** Department of Anatomy, Physiology and Genetics, Uniformed Services University of the Health Sciences, 4301 Jones Bridge Road, Bethesda, MD 20814, USA; Center for Neuroscience and Regenerative Medicine, Uniformed Services University of the Health Sciences, 4301 Jones Bridge Road, Bethesda, MD 20814, USA

**Keywords:** Traumatic brain injury, Axon damage, Neural stem cell, Intracerebroventricular, Microglia, Inflammation, Sonic hedgehog

## Abstract

Neural stem cells (NSCs) delivered intraventricularly may be therapeutic for diffuse white matter pathology after traumatic brain injury (TBI). To test this concept, NSCs isolated from adult mouse subventricular zone (SVZ) were transplanted into the lateral ventricle of adult mice at two weeks post-TBI followed by analysis at four weeks post-TBI. We examined Sonic hedgehog (Shh) signaling as a candidate mechanism by which transplanted NSCs may regulate neuroregeneration and/or neuroinflammation responses of endogenous cells. Mouse fluorescent reporter lines were generated to enable *in vivo* genetic labeling of cells actively transcribing *Shh* or *Gli1* after transplantation and/or TBI. *Gli1* transcription is an effective readout for canonical Shh signaling. In *Shh*^*CreERT2*^*;R26tdTomato* mice, *Shh* was primarily expressed in neurons and was not upregulated in reactive astrocytes or microglia after TBI. Corroborating results in *Gli1*^*CreERT2*^*;R26tdTomato* host mice demonstrated Shh signaling was not upregulated in the corpus callosum, even after TBI or NSC transplantation. Transplanted NSC expressed *Shh in vivo* but did not increase *Gli1* labeling of host SVZ cells. Importantly, NSC transplantation significantly reduced reactive astrogliosis and microglial/macrophage activation in the corpus callosum after TBI. Therefore, intraventricular NSC transplantation after TBI significantly attenuated neuroinflammation, but did not activate host Shh signaling via *Gli1* transcription.

## INTRODUCTION

Transplantation of stem cells into specific locations in the central nervous system (CNS) parenchyma may be the most appropriate approach for testing restorative cell therapies for focal lesions, such as stroke or Parkinson’s disease. However, diffuse injuries such as traumatic brain injury (TBI) may require approaches that can reach broader regions. Diffuse axonal injury in white matter tracts is the most common pathological feature of TBI [1, 2]. TBI patients often suffer long term disability [3], and no effective therapies are available to prevent the progression of white matter pathology [4, 5]. Therefore, potential therapeutics must be developed to ameliorate the progression of pathology and promote repair following diffuse axonal injury from TBI.

NSCs transplanted within the ventricular system may interact dynamically with endogenous cells to attenuate neuroinflammation, which contributes to a cascade of secondary damage in white matter tracts. Neural stem cell (NSC) transplantation also has the potential to enhance regeneration of damaged tissue directly by replacing lost cells, and/or indirectly through the synthesis of signaling factors that stimulate regenerative responses of endogenous cells in the host tissue [6]. Multipotent NSCs reside in the adult subventricular zone (SVZ) and are maintained by Sonic hedgehog (Shh) signaling [7-10]. Isolated NSCs can synthesize Shh *in vitro* after differentiation [11]. Thus, NSC transplantation may be a means to increase Shh signaling. Shh signaling and NSC transplantation have each been reported to be immunomodulatory and to promote endogenous cell repair in the corpus callosum in experimental demyelination [12-15]. However, it is presently unknown whether intraventricular NSC transplantation can modulate neuroinflammation or neuroregeneration after TBI. Additionally, approaches to demonstrate *in vivo* activation of the Shh pathway have not been used to examine this potential mechanism of NSC interaction with endogenous cells.

To address these research gaps, we used a model of experimental TBI to examine the effects of intraventricular transplantation of adult NSCs on the endogenous NSC response in the SVZ and neuroinflammation in the corpus callosum. This impact model produces traumatic axonal injury in the white matter with degenerating axons dispersed among intact axons [16-18]. The white matter pathology is similar to diffuse axonal injury in TBI patients, but is primarily in the corpus callosum over the lateral ventricles. Importantly, axon damage and neuroinflammation (reactive astrocytes and microglia/macrophages), persist in the corpus callosum out to 6 weeks post-TBI [18]. This region of pathology in the corpus callosum is adjacent to the SVZ, and thus facilitates analysis of the regenerative response of endogenous cells in the host SVZ [16, 17].

For potential clinical relevance to future autologous NSC transplantation strategies, our experimental design used intraventricular transplantation of a low dose of adult NSCs at two weeks after TBI. Analysis of the beneficial effects of NSC transplantation included both modulation of neuroinflammation and stimulation of SVZ neuroregeneration, which are each responses that can be regulated by Shh signaling. *Shh*^*CreERT2*^ and *Gli1*^*CreERT2*^ mice were each crossed to reporter lines for inducible genetic *in vivo* labeling of cells synthesizing or responding to Shh, respectively. These studies broadly examine the potential for adult NSCs transplanted into the lateral ventricles to influence endogenous cells in the adjacent white matter and SVZ regions after TBI, and specifically examine the regulatory mechanism of signaling through the Shh pathway.

## MATERIALS and METHODS

Mice were housed and cared for in accordance with the guidelines of the National Institutes of Health and the Institutional Animal Care and Use Committee of the Uniformed Services University of the Health Sciences.

### Inducible Genetic *In Vivo* Labeling of Cells Synthesizing or Responding to Shh

Mouse lines were purchased from Jackson Laboratories (Bar Harbor, ME) with information for each line listed in the supplemental materials Table S1. Mouse conditional driver lines, *Shh*^*CreERT2*^ *(Shh*^*tm2(cre/ERT2)Cjt*^*/*J), or *Gli1*^*CreERT2*^ (*Gli1*^*tm3(cre/ESR1)Alj*^*/J*), were previously shown to match mRNA transcript expression for inducible genetic labeling of cells expressing Shh [19] or cells responding to Shh signaling [20], respectively. *Shh*^*CreERT2*^ mice exhibit reporter labeling consistent with Shh expression and substantially reduce or eliminate Shh expression when crossed to mice with floxed *Shh* alleles [21]. *Gli1*^*CreERT2*^ mice exhibit reporter labeling consistent with *Gli1* expression detected by in situ hybridization [21], or using *Gli1-nLacZ* mice[22]. The *R26YFP* reporter mice (B6.129X1-Gt*(ROSA)26Sor*^*tm1(EYFP)Cos*^*/*J) and the *R26tdTomato* reporter mice (B6.Cg-Gt*(ROSA)26*^*Sortm14(CAG-tdTomato)Hze/*^J), respectively, contain the *yellow fluorescent protein* (*YFP*) or *tdTomato* fluorescent protein (*Tom*) reporter genes downstream of a floxed stop codon [23, 24]. The *R26mT/mG* reporter mice (B6.129(Cg)-Gt*(ROSA)26Sor*^*tm4(ACTB-tdTomato,-EGFP)Luo*^*/*J) ubiquitously express membrane-targeted *tdTomato* (*mT*) that switches to expression of membrane-targeted *enhanced green fluorescent protein* (*mG*) after Cre-mediated recombination [25]. *Gli1*^*CreERT2*^ mice were crossed to either *R26YFP* or *R26tdTomato* mice. *Shh*^*CreERT2*^ were crossed to either *R26tdTomato* or *R26mT/mG* mice. First-generation crosses were used for experimental procedures. To induce nuclear translocation for Cre-mediated recombination, 10 mg of tamoxifen (TMX; Sigma, St. Louis, MO) in corn oil was administered by oral gavage daily on days 2 and 3 after injury or NSC transplant [17].

### Generation and Characterization of Adult NSCs

NSCs were isolated from the SVZ of 8-10 week old mice and maintained as neurospheres, as in previously published protocols [26, 27]. For NSC isolation, the dissected brain was cut into 1 mm coronal slices. The SVZ was dissected and enzymatically digested using the Neural Tissue Dissociation Kit (P) (Miltenyi Biotec Inc., San Diego, CA), followed by mechanical dissociation. Cells isolated from each mouse were plated in 10 ml of mouse NeuroCult NSC Basal Medium (StemCell Technologies, Vancouver, BC, Canada) supplemented with mouse NeuroCult NSC Proliferation Supplement (StemCell Technologies), 20 ng/ml rmEGF (Life Technologies, Grand Island, NY City, State), 10 ng/ml of rhFGF-b (Life Technologies), 2 μg/ml Heparin (Life Technologies) and 100 U/ml Pen/Step (Life Technologies). Cells were cultured at 37°C with 5% CO_2_. Following the first passage, cells were dissociated to a single cell suspension in 0.025% Trypsin-EDTA (Invitrogen, Carlsbad, CA) and reseeded in culture medium at 1 × 10^5^ cells/ml. Neurospheres from each isolation were cryopreserved at passage two in F12/DMEM (ThermoFisher) with 15% dimethylsulfoxide (DMSO, ATCC). Cells were resurrected at least 2 weeks prior to transplantation and passaged at least once, but no more than five times, prior to transplantation (Fig. S1). NSCs with constitutive green fluorescent protein (GFP) labeling (NSC-GFP) were generated from C57BL/6-*Tg(UBC-GFP)30Scha/*J) mice. *Shh*^*CreERT2*^*;R26mT/mG* mice were used to generate NSC-Shh cells. After TMX administration, cells with constitutive mT labeling convert to mG labeling if the *Shh* promoter is transcriptionally active *in vivo*.

To quantify the number of cells in primary neurosphere cultures, an aliquot of cells was taken from a single cell suspension of NSCs following extraction from a single mouse and after each passage. The aliquot of cells was diluted 1:1 in 0.4% trypan blue (Sigma) to determine cell viability based on trypan blue exclusion. The total number of cells (live + dead cells) were quantified using a phase contrast microscope and a hemocytometer. Starting at the first passage, the growth rate of cells was determined by dividing the total number of cells at the time of passage by the number of cells plated. The growth rate was then used to determine the cumulative number of cells following each passage to generate a growth curve. Proliferation through passage eight followed a linear growth curve with log phase amplification that is characteristic of healthy stem cell self-renewal (Fig. 1A).

**Figure 1.**
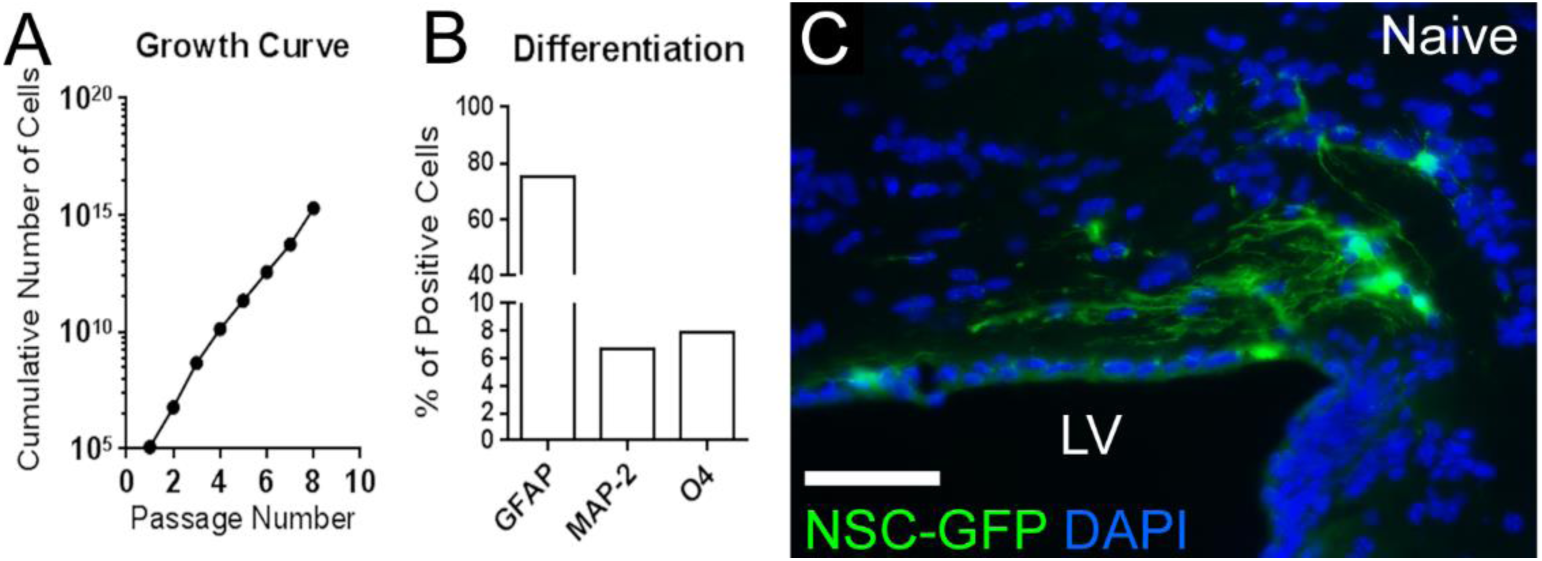
Adult NSC *in vitro* characterization and NSC-host response after transplantation. **A:** *In vitro* growth curve of cumulative cells generated demonstrates proliferation. **B:** *In vitro* confirmation of multipotency by quantification of differentiated cells immunolabeled for distinct lineage markers: astrocytes with GFAP, neurons with MAP-2, or oligodendrocytes with O4. **C:** NSCs ubiquitously expressing green fluorescent protein (NSC-GFP) formed processes 1 week following transplantation into the parenchyma adjacent to the lateral ventricle (LV). Sections were counterstained with 4′,6-diamidino-2-phenylindole (DAPI, blue). Scale bar = 50 μm (C).

NSCs were plated at densities of 4.8 x 10^4^ cells/ml for clonal analysis or 80,000 cells per coverslip and characterized as in previously published protocols [14]. Cultures were immunolabeled for markers of differentiation along the astroglial lineage (glial fibrillary acidic protein, GFAP; rabbit polyclonal, 1:100; DAKO, Carpinteria, CA), the neuronal lineage (microtubule associated protein-2, MAP-2; rabbit polyclonal, 1:200; Abcam, Cambridge, MA), and the oligodendroglial lineage O4 (mouse monoclonal, 1:10; [28]). Secondary antibodies used were goat anti-rabbit IgG conjugated with Alexa Fluor 488 (Life Technologies,) to detect GFAP and MAP-2, and goat anti-mouse IgM conjugated with FITC (Jackson ImmunoResearch, West Grove, PA) to detect O4. Neurospheres dissociated after passage five and cultured in differentiation medium demonstrated multipotency based on differentiation into astrocytes (GFAP), neurons (MAP-2), and oligodendrocyte lineage cells (O4) (Figs. 1B, S1).

Adult NSCs isolated from mice ubiquitously expressing GFP (NSC-GFP) (Fig. S1), and transplanted into naïve mice, demonstrated *in vivo* viability based on integration and extension of processes within the corpus callosum (Fig. 1C).

### Lentiviral Transduction of Human non-NSCs

Lentiviral transduction was used to generate human embryonic kidney 293 (HEK293) cells expressing tdTomato (HEK-Tom). The pLVX-IRES-tdTomato vector (Clontech Laboratories, Mountain View, CA) was co-transfected with the viral core packaging construct PAX2 and the VSV-G envelope protein vector pCAG VSV-G into HEK293T cells (ATCC CRL-1573; Manassas, VA) to generate lentivirus for expression of tdTomato (Lenti-Tom), using previously published protocols [29]. Approximately 70% of the Lenti-Tom transduced HEK293T cells expressed the tdTomato reporter. HEK293 cells were authenticated as human by short tandem repeat profiling of an aliquot of the transfected cells using a 16 marker interspecies contamination test that resulted in > 80% match to human (IDEXX BioResearch, Columbia, MO(ATCC, Manassas, VA).

### Surgical Procedures for TBI

TBI was produced in male (8–10 week old) *Gli1*^*CreERT2*^*;R26tdTomato, Shh*^*CreERT2*^*;R26tdTomato* or C57BL/6J mice, as previously detailed [16, 17]. Briefly, an Impact One(tm) Stereotaxic Impactor device was aligned with a 3-mm flat tip to impact the skull at bregma using settings of 1.5 mm depth, 4.0 m/s velocity (video recorded as 4.7 m/s) and 100ms dwell time. These parameters have been shown to produce pathology in the corpus callosum with axon damage, astrogliosis, and microglial activation, similar to our previously characterized results [18]. Mice were excluded if the impact resulted in apnea > 30 seconds or depressed fracture. Post-surgical data was recorded to characterize the injury across mouse lines. The righting reflex was significantly increased after TBI, compared to sham, in C57BL/6J mice (sham 0.56 ± 0.12, n = 5; TBI 5.2 ± 0.58, n = 5; p < 0.0001), *Shh*^*CreERT2*^*;R26tdTomato* mice (sham 1.90 ± 0.60, n = 5; TBI 7.20 ± 1.91, n = 10; p = 0.0101), and *Gli1*^*CreERT2*^*;R26tdTomato* (sham 2.11 ± 0.31, n = 9; TBI 5.43 ± 0.719, n = 8; p = 0.0004). The righting reflex after TBI was not affected by the genetic background of these mouse lines (p = 0.3666). Surgical data includes only mice used for quantitative analysis in subsequent experiments. Fractures on the skull or contusions on the surface of the brain were not observed at the time of dissection. Sham surgery included anesthesia and scalp incision. After surgery, mice were randomly selected for vehicle or cell transplantation.

Two weeks prior to HEK-Tom cell transplantation, a cohort of mice received a controlled cortical impact (CCI). The CCI model of TBI was used to examine a more invasive injury with focal gray matter damage and blood-brain barrier disruption, as previously detailed [17]. Briefly, in male (8–10 week old) *Gli1*^*CreERT2*^*;R26YFP* or C57BL/6J mice, a craniotomy was performed to expose the dura mater. An Impact One(tm) Stereotaxic Impactor device (Leica Biosystems; Buffalo Grove, IL) was used to impact the dura mater of the cerebral cortex of the right hemisphere (0 bregma; 1.5 mm lateral). Settings used were 2 mm tip diameter, 1 mm depth, 1.5 m/s velocity, and 100 ms dwell time. Mice used for CCI were *Gli1*^*CreERT2*^*;R26YFP* (sham n = 1; CCI n = 3).

### Cell Transplantation into the Lateral Ventricle

Host mice for transplantation experiments were C57BL/6J, *Shh*^*CreERT2*^*;R26tdTomato, Gli1*^*CreERT2*^*;R26tdTomato, or Gli1*^*CreERT2*^*;R26YFP*. Briefly, a burr hole of approximately 1.0 mm diameter was drilled into the skull, unless a craniotomy for CCI was previously performed. A 10 μl Hamilton gas tight syringe (Cat# 7653-01; Hamilton Company, Reno, NV) with compression fitting adapters (Cat# 55750-01; Hamilton Company) and a pulled glass micropipette (outer diameter 50 μm) was used to microinject 2 μl of cell suspension (2.5 X 10^4^ cell/ul) or F12/DMEM vehicle (ThermoFisher) into the lateral ventricle (1.0 mm to the right of bregma and -1.75 mm deep). Transplanted cells were either NSC-GFP cells, NSC-Shh cells or HEK-Tom cells.

### Immunohistochemistry in Tissue Sections

Mice were perfused with 3% paraformaldehyde and brains were processed as in previously published protocols ([17]. Immunohistochemistry was performed to detect astrocytes expressing GFAP (rabbit polyclonal, 1:1000; DAKO), microglia/macrophage with CD11b (rat monoclonal, 1:100; AbCam,), or T-lymphocytes with the pan T-cell marker CD3 (rabbit polyclonal, 1:2000; DAKO,). Neurons were immunolabeled with neuronal nuclei marker NeuN (mouse monoclonal, 1:100; Sigma). Axons were identified by localization of neurofilaments (rabbit polyclonal neurofilament H; 1:500; Encor Biotechnology, Gainesville, FL). Immunolabeling of nonphosphorylated neurofilaments (mouse monoclonal SMI-32; 1:1000; Covance, Gaithersburg, MD) was used to detect damaged axons [30]. YFP reporter expression was detected with an antibody against GFP (rabbit polyclonal, 1:1000; Life Technologies). In *Gli1*^*CreERT2*^*;R26tdTomato* or *Gli1*^*CreERT2*^*;R26YFP* mice, goat anti-rabbit IgG conjugated with Alexa Fluor 488 (Life Technologies) was used to detect GFAP, CD3 or YFP, and goat anti-rat IgG conjugated with Alexa Fluor 488 (Life Technologies) was used to detect CD11b. In C57BL/6J or *Gli1*^*CreERT2*^*;R26tdTomato* mice, goat anti-rat IgG conjugated with Alexa Fluor 555 (Life Technologies) was used to detect CD11b or donkey anti-rabbit IgG conjugated to Cy3 (Jackson Immunoresearch) was used to detect GFAP. Sections were counterstained with 4′,6′-diamidino-2-phenylindole (DAPI; Sigma) then mounted with Vectashield (Vector Laboratories).

### Quantification and Statistical Analysis

The number of mice used from each line is noted with the TBI surgical methods. Conditions were randomly assigned within each mouse cohort. All quantification was performed blinded to injury (TBI/sham) and transplantation (cells/vehicle) procedures. All tissue analysis was performed in coronal brain tissue sections. Analyses of the corpus callosum or SVZ used at least three sections per mouse distributed between approximately + 0.3 mm to - 0.2 mm relative to bregma. Analyses of NSC-Shh cells within the lateral and third ventricles used sections from between approximately - 0.0 mm to -1.6 mm relative to bregma, which were sampled at least every 300 μm to reach a minimum of 50 NSC-Shh cells counted in each mouse. Images were collected using Spot RT3 digital camera (Diagnostic Instruments, Sterling Heights, MI). Spot Advanced software was used to measure the area of the corpus callosum and SVZ, as previously described [16, 17]. The area of GFAP or CD11b immunolabeling in the corpus callosum was quantified using Metamorph software (Molecular Devices, Downingtown, PA) to threshold pixels with intensities above background levels as previously detailed [16, 18]. CD11b cells within the corpus callosum were also counted individually to differentiate activation stage as resting (fine processes), activated (hypertrophic with thickened processes), or ameboid (retracted processes) as previously characterized [16].

Prism 6.0 (GraphPad Software) was used for graphing and statistical analysis. Two-way analysis of variance (ANOVA) was performed to determine significant differences between NSC transplanted and vehicle injected cohorts across naïve, sham, and/or injured conditions. Significant differences between the righting response in sham and TBI cohorts of a single genotype were determined using the two tailed unpaired *t*-test, while one way ANOVA was used to compare TBI conditions across the three genotypes. Values shown are mean ± SEM. Statistical significance was determined as *p* < 0.05.

## RESULTS

### Induced Genetic *In Vivo* Labeling of Host Cells Expressing Shh

An impact model of TBI was used that produces axon damage in the corpus callosum, primarily above the lateral ventricles [16-18]. This injury localization facilitates analysis of potential effects from NSC cells transplanted into the adjacent ventricle in subsequent experiments. We first examined *Shh* expression following TBI using TMX administration to induce genetic labeling of cells actively transcribing *Shh in vivo* based on expression of the tdTomato reporter (Shh-Tom) in *Shh*^*CreERT2*^*;R26tdTomato* mice (Fig. 2A, B). In both TBI and sham mice, TMX administered after surgery induced Shh-Tom labeling predominantly in neuronal cell bodies, axons, and processes based on morphology and immunolabeling for NeuN (Figs. 2C-I, S2). Shh-Tom neurons were distributed as previously characterized in adult mouse brain [31, 32]. Shh-Tom neurons were present in the striatum, adjacent to the SVZ and extending processes to the SVZ (Fig. 2C), and in the cerebral cortex, including regions under the site of impact (Fig. 2D, E). Astrocytes, identified by expression of GFAP, did not co-label for Shh-Tom in the cortex (2D, E) or in the corpus callosum, even after TBI in regions with reactive astrocytes (Fig. 2F, G). Indeed, Shh-Tom labeling of cell bodies was very rarely observed in the corpus callosum. Shh-Tom labeled axons could be followed traversing the corpus callosum in both sham (F, H) and TBI (G, I) mice. Rare evidence of damage, i.e. swelling and beading, in Shh-Tom labeled axons was observed only after TBI (Fig. 2I). These findings indicate that TBI does not stimulate Shh synthesis in reactive astrocytes and that the overall pattern of cells expressing Shh is similar between sham and TBI mice.

**Figure 2.**
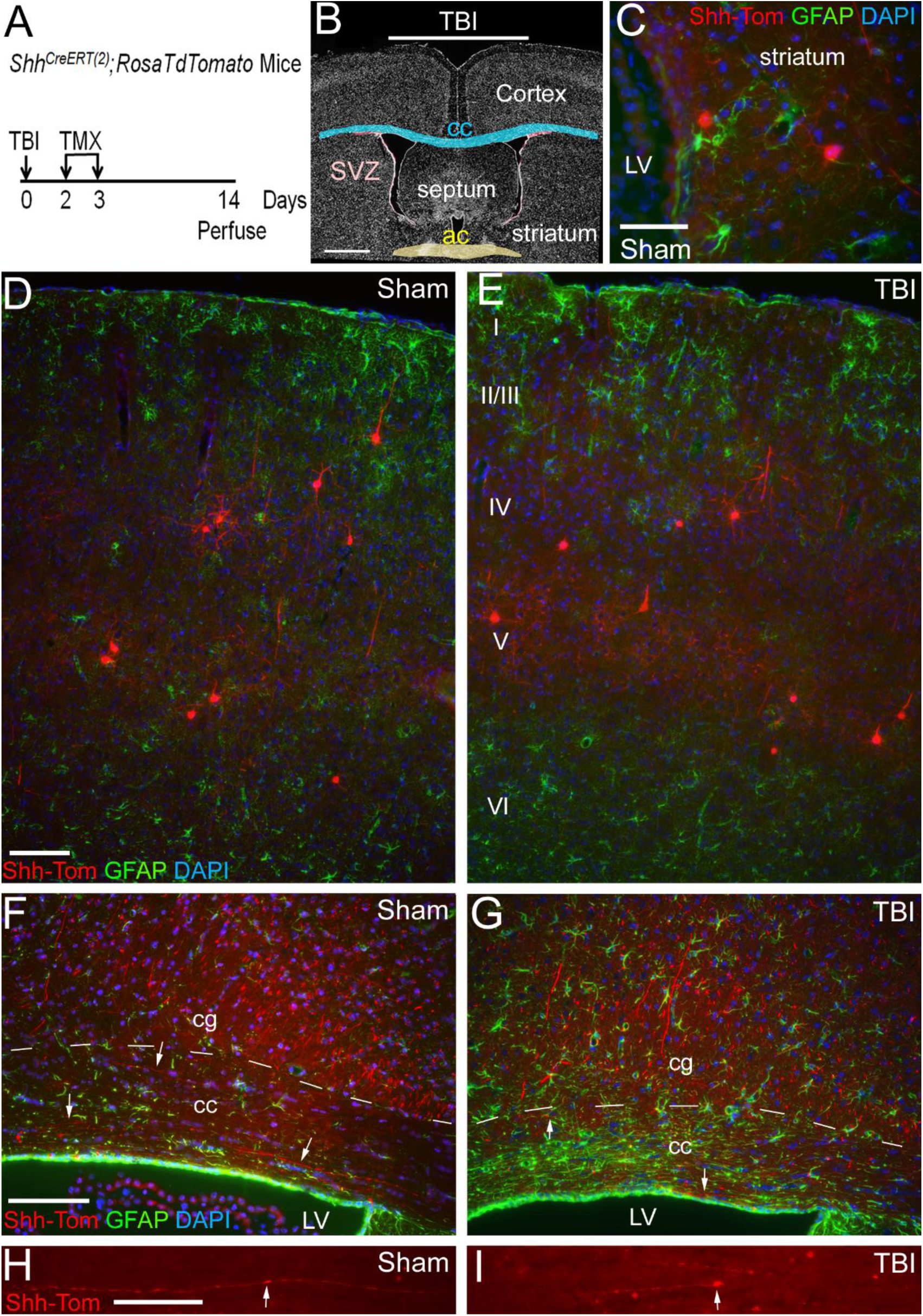
Induced genetic *in vivo* labeling to detect *Shh* expression following traumatic brain injury (TBI). **A:** *Shh*^*CreERT2*^*;R26tdTomato* mice received tamoxifen (TMX) 2 and 3 days after TBI or sham surgery and were perfused at 2 weeks post-surgery. **B:** Diagram showing impact to the skull at bregma corresponded with the coronal level of the anterior commissure (ac). **C-G:** Coronal sections with *Shh* transcription evident via the tdTomato reporter (Shh-Tom, red). GFAP immunolabeling (green) to detect astrocytic response to TBI. DAPI nuclear stain (blue) to identify cortical layers (I-VI). **C:** Shh-Tom cells with neuronal morphology in the striatum near the SVZ and lateral ventricle (LV). Similar Shh-Tom cells in the striatum are colabeled with the neuronal marker NeuN (Fig. S2). **D-E:** Shh-Tom labeling in distributed cortical cells identified as neurons by morphology (D, E) and co-labeling with NeuN (Fig. S2). GFAP immunolabeling in the cortex showed mild cortical astrogliosis after TBI (D, E). Shh-Tom cells did not have an astrocytic morphology or co-label for GFAP (D, E). **F-G:** Shh-Tom labeling of axons in the cingulum (cg) and corpus callosum (cc) was similar in sham (F) and TBI (G) mice. After TBI, reactive astrocytes in the corpus callosum were not co-labeled with Shh-Tom (G). **H-I:** Shh-Tom axons were found in the corpus callosum but at a very low density. Normal variation of Shh-Tom labeling along the length of axons in a sham mouse (arrow; H). Consistent with axon damage after TBI, rare pathological swellings were observed in Shh-Tom labeled axons (arrow; I). Scale bars = 1 mm (B), 100 μm (C, D, F), 50 μm (H). Representative images are shown from analysis of a cohort of sham (n = 3) and TBI (n = 4) mice.

We next explored the potential for NSC transplantation to stimulate Shh signaling after TBI. *Shh*^*CreERT2*^*;R26tdTomato* host mice had TBI or sham surgery and 2 weeks later received NSC-GFP cell transplant, or a vehicle injection, followed by perfusion for tissue analysis at 4 weeks post-TBI or sham surgery (Fig. 3A, B). Shh-Tom labeling near the lateral ventricles was similar to the labeling observed at two weeks after sham or TBI surgery (Fig. 2C, 3C, S2B). Primarily, Shh-Tom labeling was observed in cells with neuronal morphology, including cells in the striatum near the SVZ and axons extending toward the ventricle from septal regions (Fig. 3C). Shh-Tom labeling was not altered by the close proximity of transplanted NSC-GFP cells (Fig. 3D-H). Furthermore, with and without NSC-GFP transplantation, Shh-Tom labeling of axons appeared similar even in the corpus callosum and cingulum, which are regions with axon damage in this TBI model (Fig. S3). Finally, near the injection tract, reactive astrocytes did not exhibit Shh-Tom labeling (data not shown). Overall, neither TBI nor NSC-GFP transplantation markedly changed the pattern of host cells expressing Shh.

**Figure 3.**
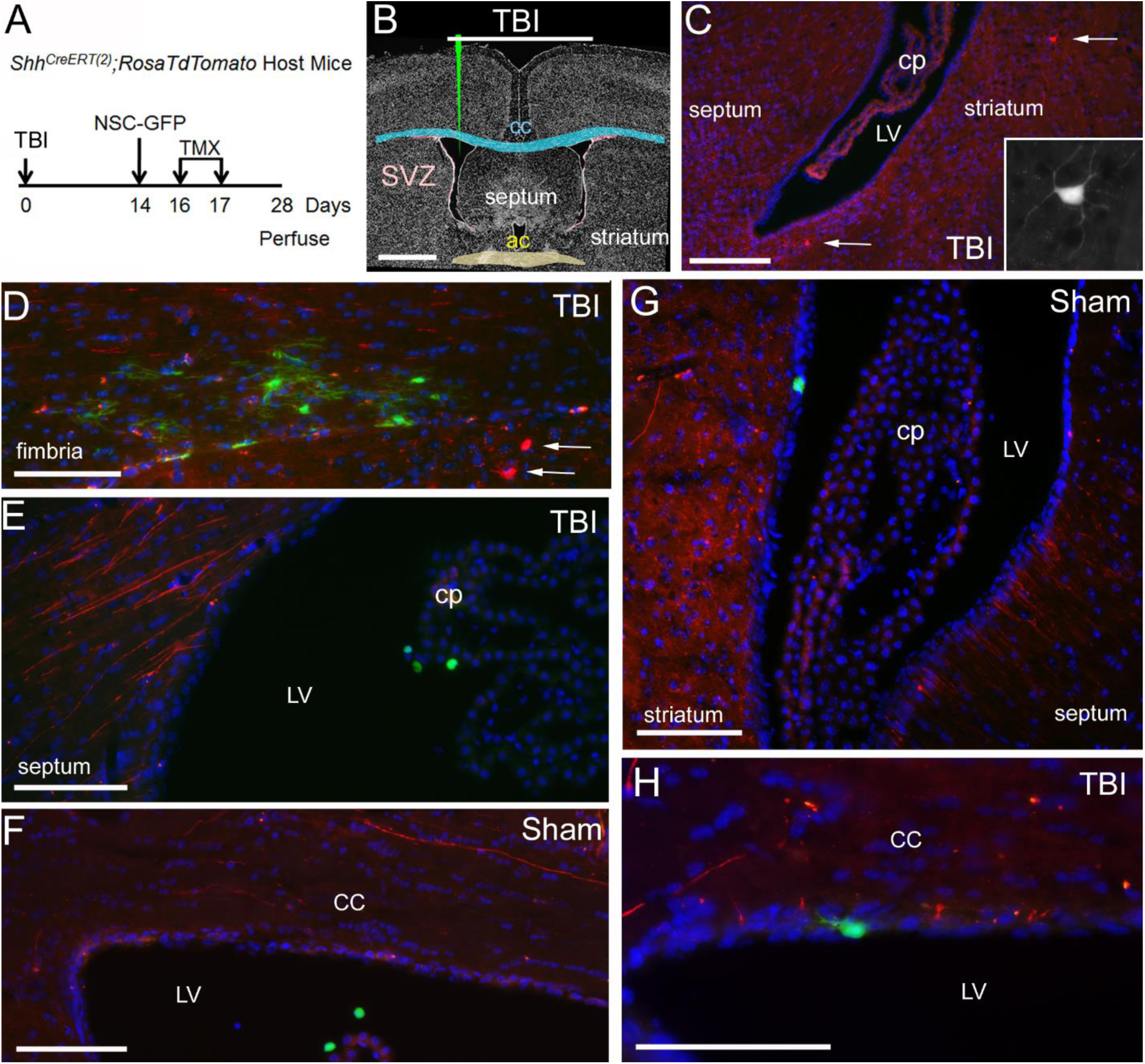
Induced genetic *in vivo* labeling of host cells expressing *Shh* after NSC transplantation into mice with traumatic brain injury (TBI). **A, B:** Timeline (A) and diagram (B)for experiments in *Shh*^*CreERT2*^*;R26tdTomato* host mice. Two weeks after TBI or sham surgery mice received an injection into the lateral ventricle (LV) of either NSCs constitutively expressing green fluorescent protein (NSC-GFP cells), or vehicle. To induce genetic labeling of host cells expressing *Shh* (Shh-Tom cells), TMX was administered on days 2 and 3 after injection of NSCs/vehicle. Mice were perfused 4 weeks after TBI/sham surgery. **C-H:** Coronal sections showing Shh-Tom labeling (red) with DAPI nuclear stain (blue) and transplanted NSC-GFP cells (green). Shh-Tom labeling in cell bodies (C-D arrows) and axons (E-H). Inset (C) shows neuronal morphology of Shh-Tom cells (arrows) in the striatum with example from upper arrow. Injections occasionally missed the LV resulting in NCS-GFP transplantation into the adjacent tissue and serve to show the potential to elaborate extensive processes is dependent on the transplant site (D). NSC-GFP cells transplanted directly into the LV grew as spheres (E, F), and were often observed nestled in the choroid plexus (cp), or extended only short processes when adhered to the walls of the LV (G, H). Scale bars = 1 mm (B), 200 μm (C), 100 μm (D-G), 50 μm (H). Representative images are shown from analysis of a cohort of sham + NSC-GFP (n = 2), TBI + vehicle (n = 3), and TBI+ NSC-GFP (n = 3) mice.

### Induced Genetic *In Vivo* Labeling Demonstrates NSCs Express Shh After Transplantation

NSCs have been shown to synthesize Shh following *in vitro* differentiation [11]. Therefore, we examined the potential for NSCs to express *Shh* after transplantation. SVZ NSCs were cultured from *Shh*^*CreERT2*^*;R26mT/mG* mice (NSC-Shh cells) to be able to detect transplanted NSCs with and without *Shh* expression *in vivo*. In NSC-Shh cells actively transcribing *Shh*, TMX administration induces a switch of membrane labeling from tdTomato (red) to GFP (green). Overlap appears as yellow in cells that had pre-existing membranes labeled by tdTomato and then have begun to express GFP. Two weeks post-TBI or sham surgery, NSC-Shh cells were transplanted into the lateral ventricle of C57BL/6J mice and TMX was delivered on days 2 and 3 after transplantation (Fig. 4A, B). After injection into the lateral ventricle, NSC-Shh cells distributed in both the lateral ventricles and the third ventricle, which were analyzed as sites near and remote from the injury, respectively (Fig. 4C). TMX induced GFP expression indicative of active *Shh* expression *in vivo* (Fig. 4D). In sham mice, approximately 60-80% of the transplanted NSC-Shh cells exhibited GFP labeling in either the lateral ventricle or the third ventricle (Fig. 4D, E-E”). Remarkably, TBI resulted in a significant reduction in the proportion of NSC-Shh cells labeled with GFP (Fig. 4D, F-F”). This suppression of *Shh* expression in NSC-Shh cells after TBI was only found among cells in the lateral ventricle region, indicating a specific NSC response based on proximity to the injury (Fig. 4D).

**Figure 4.**
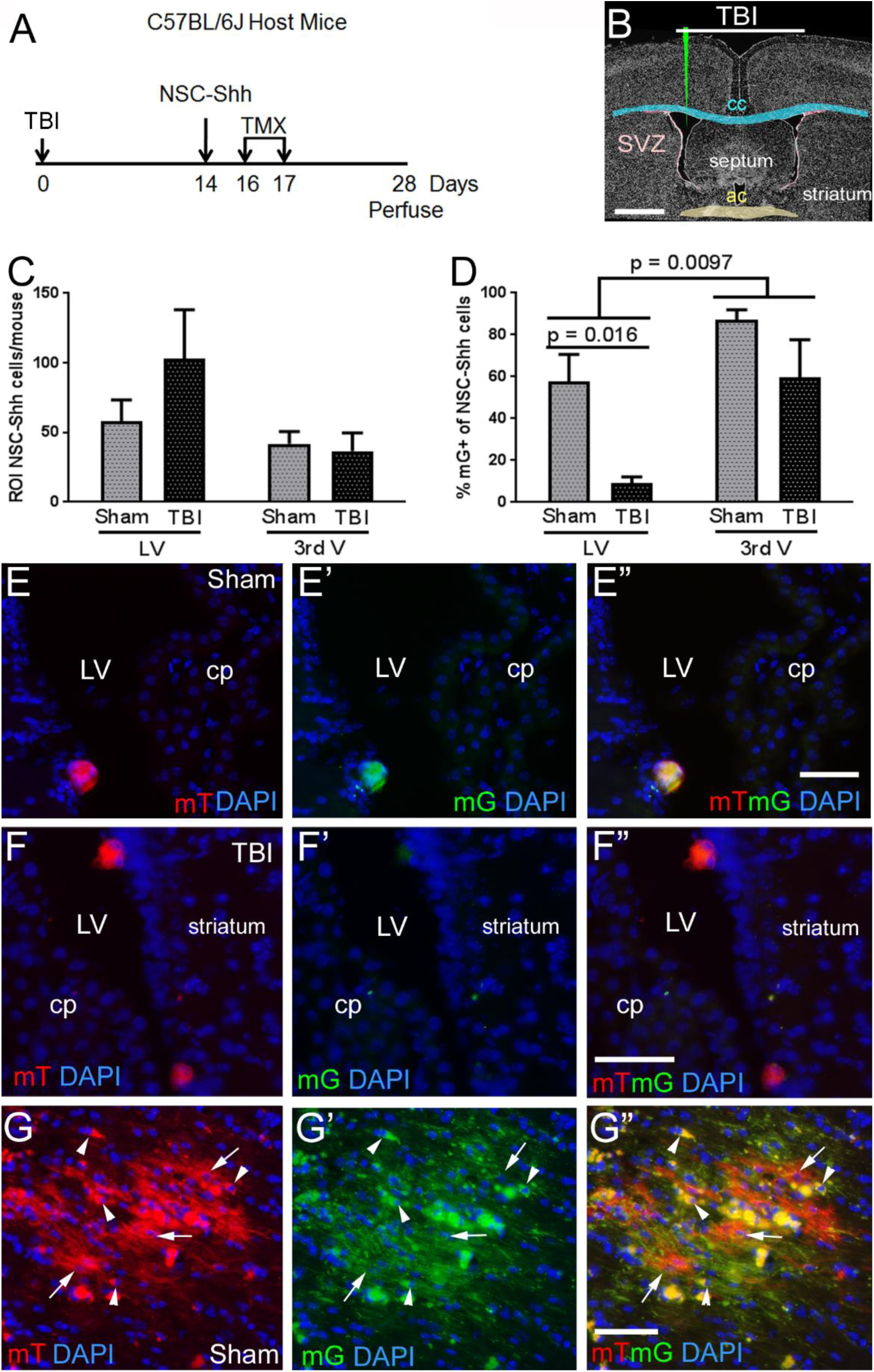
Induced genetic *in vivo* labeling of NSCs expressing *Shh* after transplantation. **A-B:** Timeline (A) and diagram (B) for host C57BL/6J mice that had TBI or sham surgery and 2 weeks later received an injection into the lateral ventricle (LV) of NSCs isolated from *Shh*^*CreERT2*^*;R26mT/mG* mice (NSC-Shh cells). Tamoxifen (TMX) was administered to induce a genetic switch from constitutive mT (red) to mG (green) fluorescence in transplanted NSC-Shh cells expressing *Shh*. Membranes that were already labeled by constitutive mT and then begin to express mG after TMX may exhibit both mT and mG, so that co-labeled membranes appear yellow. Mice were perfused at 4 weeks after TBI or sham surgery. **C:** Transplanted NSC-Shh cell distribution within the lateral ventricles and third ventricle. **D:** Proportion of transplanted NSC-Shh cells that exhibited expression of mG, indicating *in vivo Shh* expression. In TBI mice, mG labeling was significantly reduced near the site of injury in the LV but not in the third ventricle. **E-F”:** Coronal sections showing transplanted NSC-Shh cells and DAPI nuclear stain (blue) to visualize the choroid plexus (cp) and wall of the LV. Sham mice shown with NSC-Shh cells in the LV that expressed both the mT (red) and mG (green) reporters, with overlap (yellow), indicating *in vivo* expression of *Shh* after transplantation (E-E”). TBI mice shown with two clusters of NSC-Shh cells in the LV that express constitutive mT (red) but not *Shh* driven mG (green) (F-F”). *Shh* driven mG expression was more evident in cells transplanted into the parenchyma, which extended processes (G-G”, fimbria). After TMX induction, membranes appear green in newly elaborated processes, but appear yellow in cell bodies where mT and mG overlap (arrowheads). Other transplanted cells and processes are labeled by only mT expression (arrows), indicating lack of *Shh* expression or inefficiency of Cre recombination *in vivo*. Scale bars = 1 mm (B), 50 μm (D, F), 25 μm (E). ROI = region of interest quantified for designated rostrocaudal extents of the lateral ventricles and the third ventricle (see Methods). Quantification included analysis of a cohort of sham (n = 5) and TBI (n = 5) mice.

The majority of NSC-Shh cells in the ventricles had a rounded immature morphology (Fig. 4E-F”). For comparison with the intraventricular environment, NSC-Shh cells are shown after transplantation directly into white matter adjacent to the ventricles (Fig. 4G-G”). NSC-Shh cells in the parenchyma often elaborated processes and were labeled with GFP, indicating active *Shh* expression *in vivo* (Fig. 4G-G”). In NSC-Shh cells that did not express *Shh*, the membranes of the cell bodies and extended processes continue to be labeled with tdTomato. In NSC-Shh cells that expressed *Shh in vivo,* the switch to *GFP* expression results in only GFP labeling of new membranes synthesized as cell processes form. In contrast, cell bodies appear yellow from the overlap of pre-existing tdTomato labeled membrane and the subsequent GFP membrane labeling.

### Analysis of NSC Transplantation on Shh Signaling in Host SVZ Cells

Shh signaling regulates NSC maintenance in the SVZ of adult mice [10, 21]. Therefore, we next examined whether NSCs transplanted into the lateral ventricle would increase Shh signaling in endogenous cells in the adjacent SVZ after TBI (Fig. 5A-C). NSC-GFP cells (Fig. S1) were transplanted into *Gli1*^*CreERT2*^*; R26tdTomato* mice. TMX was administered on days 2 and 3 post-transplantation to detect changes in the population of endogenous cells responding to Shh signaling, based on induced genetic labeling with tdTomato (Glil-Tom). Within the lateral ventricle, NSC-GFP cells were typically adhered to the walls of the ventricle and nestled within the choroid plexus (Figs. 5, S4). In comparison with naïve mice (Fig. 5D), the endogenous Gli1-Tom population in the SVZ was not altered by the microinjection technique or NSC-GFP transplantation in either sham mice (Fig. 5E) or after TBI (Fig. 5F). The overall pattern of Gli1-Tom labeling in the areas adjacent to the lateral ventricles was also similar across conditions after vehicle or NSC-GFP injection (Fig. S4). In addition to the SVZ, Gli1-Tom labeling was observed in distributed cells in the choroid plexus within the lateral ventricles (Figs. 5, S4). Gli1-Tom cells were only very rarely observed in the corpus callosum, even after TBI (Figs. 5, S4).

**Figure 5.**
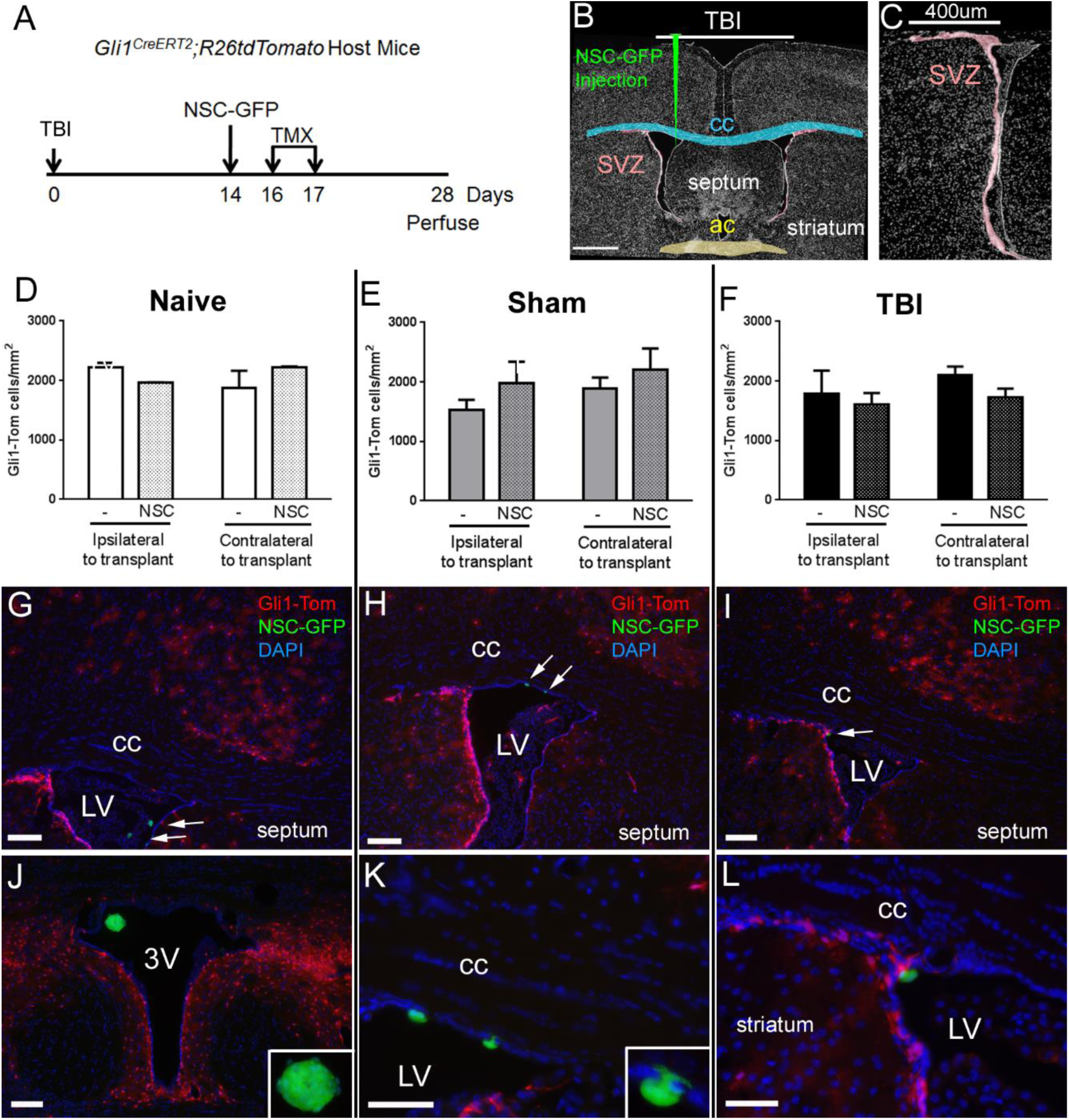
Induced genetic *in vivo* labeling of host cells expressing *Gli1* in the subventricular zone (SVZ) following traumatic brain injury (TBI) and NSC transplantation. **A-C:** Timeline for *Gli1*^*CreERT2*^*;R26tdTomato* host mice (A) with diagrams of TBI and injection site (B) and the SVZ region quantified (C) in naïve, sham, or TBI conditions. NSC-GFP cells for transplantation were isolated from mice constitutively expressing green fluorescent protein. **D-F:** Quantification of cells actively responding to canonical Shh signaling (Gli1-Tom cells), in the SVZ showed no significant change in naïve (D), sham (E), or TBI (F) mice following injection of NSC-GFP as compared to vehicle. Low magnification images of the full SVZ regions is provided in Fig. S4. **G-L:** NSC-GFP cells transplanted into the lateral ventricle (LV) nestled within the choroid plexus (G) or adhered to the walls of the lateral ventricles (H, I) and were also found within the third ventricle (J). Gli1-Tom labeled cells are not evident in the corpus callosum (cc). In contrast, robust Gli1-Tom labels cells are found in the SVZ and in the cortex. Higher magnification of cells shown at the arrows (H, I). One cell has processes extended around the ependymal cell layer (K, inset). Scale bars = 1 mm (B), 400 μm (C), 200 μm (G-J), 50 μm (K, L). Quantification included cohorts of naïve + vehicle (n = 3), naïve + NSC-GFP (n = 2), sham + vehicle (n = 4), sham + NSC-GFP (n = 4), TBI + vehicle (n = 3), and TBI + NSC-GFP (n = 4).

### Analysis of NSC Transplantation on Neuroinflammation and Shh Signaling in the Corpus Callosum

We next evaluated the effect of NSC transplantation on TBI-induced pathology (Fig. 6). This TBI model produces well characterized axonal damage and neuroinflammation in the corpus callosum [16, 18]. As shown in these prior studies, axon damage at longer times points after injury may not be adequately quantified based on β-amyloid precursor protein accumulation. Therefore, axon damage was detected using SMI-32 antibody, which recognizes non-phosphorylated neurofilaments [30]. Analysis of damaged axons focused on the cingulum, under the impact site, to image axons in cross-section for more accurate quantification (Fig. 6B-D). This analysis was performed using the *Gli1*^*CreERT2*^*;R26tdTomato* as analyzed in Fig. 5. TBI significantly increased axon damage when compared to sham mice. However, NSC transplantation did not have a significant effect on reducing axon damage.

**Figure 6.**
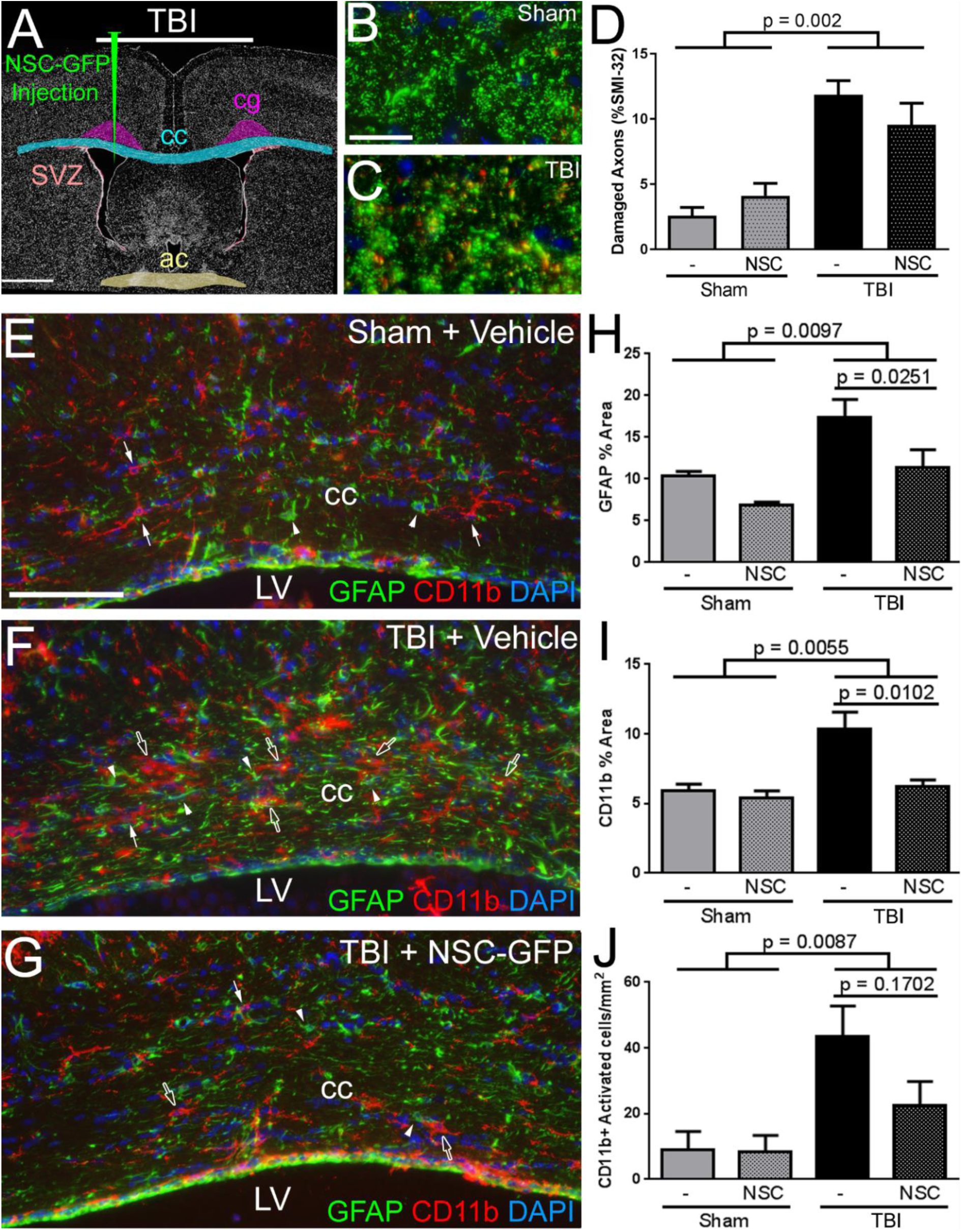
NSC transplantation into the lateral ventricle (LV) following traumatic brain injury (TBI) suppresses injury-induced neuroinflammation in the corpus callosum. **A:** Experimental diagram showing corpus callosum (cc) and cingulum (cg) regions quantified at the coronal level of the anterior commissure (ac) for mice prepared as detailed in Figs. 5 and S4. Mice were analyzed at two weeks after NSC/vehicle injection which was four weeks after TBI/sham surgery. **B-D:** Axons in the cingulum cut through transversely exhibit immunoreactivity for total neurofilaments (green) or SMI-32 (red) to detect non-phosphorylated neurofilaments as an indicator of axon damage. SMI-32 labeled axons are significantly increased in TBI, as compared to sham mice (D). **E-G:** Coronal sections of the corpus callosum immunostained for GFAP to detect astrocytes (green, arrowheads) and Cd11b to identify microglia/macrophages (red, arrows) with DAPI nuclear stain (blue). As compared to sham mice (E), immunoreactivity for reactive changes in GFAP and CDllb are evident after TBI (F) and attenuated in TBI mice with NSC transplantation (G). **H-J:** Quantification of neuroinflammation in the corpus callosum. Immunoreactivity for GFAP (H) and CD11b (I, J) was significantly increased in mice with TBI. NSC-GFP transplantation following TBI significantly reduced GFAP (H) and CD11b (I) immunoreactivity. Scale bar = 1 mm (A), 20 μm (B), 100 μm (E). Quantification included cohorts of sham + vehicle (n = 3), sham + NSC-GFP (n = 3), TBI + vehicle (n = 3), TBI + NSC-GFP (n = 3 for neurofilaments; n = 4 for CDllb GFAP).

Shh signaling may also modulate neuroinflammation, i.e. astrocyte and microglial reactivity, in the corpus callosum, as has been reported after experimental demyelination [13]. Therefore, we examined the potential for transplanted NSC-GFP cells to attenuate TBI-induced neuroinflammation in the corpus callosum adjacent to the lateral ventricles (Fig. 6E-J). TBI resulted in astrogliosis and macrophage/microglia activation in the corpus callosum that persisted at 4 weeks post-TBI (Fig. 6F, G), as compared to sham mice (Fig. 6E). Quantification of GFAP immunoreactivity showed an increase in astrocyte reactivity after TBI that was significantly reduced by NSC-GFP transplantation (Fig. 6H). Quantification of CD11b immunoreactivity also demonstrated increased activation of microglia/macrophages after TBI that was significantly reduced by NSC-GFP transplantation (Fig. 6I). Furthermore, CD11b immunolabeled cells with an activated morphology (hypertrophic processes) were also increased after TBI and reduced by NSC-GFP transplantation (Fig. 6J). These results demonstrate that NSC-GFP transplantation suppressed white matter neuroinflammation induced by TBI (Fig. 6H-J).

Only very rare Gli1-Tom labeled cells were detected in the corpus callosum of these *Gli1*^*CreERT2*^*;R26tdTomato* mice (Fig. 5D-I). This result demonstrates that the TBI pathology did not activate canonical Shh signaling in reactive astrocytes or microglia. Furthermore, the immunomodulatory effect observed from NSC transplantation is likely mediated through pathways that do not activate canonical Shh signaling.

### Activation of Shh Responsive Cells in Host SVZ After Invasive Brain Injury and HEK Cell Transplantation

To more accurately interpret the lack of changes in Gli1-Tom labeling among endogenous cells after TBI and/or NSC transplantation, a further set of experiments was designed as a positive control to elicit induced genetic *in vivo* labeling of Shh responsive cells in the SVZ. Shh signaling has been shown to increase after invasive injury, which involves a robust neuroinflammatory response [17, 33]. Therefore, we predicted that a strong immune rejection response would be observed after transplantation of HEK cells, which are human non-NSC cells. And, using controlled cortical impact (CCI), which is a more invasive form of TBI, would be more permissive to an immune response due to cortical tissue damage and breakdown of the blood-brain barrier. At 2 weeks after CCI, HEK cells expressing the tdTomato reporter (HEK-Tom cells) were transplanted into *Gli1*^*CreERT2*^*:R26YFP* mice (Fig. 7A). Mice were administered TMX on days 2 and 3 after the HEK-Tom cell transplantation. Immunohistochemistry for CD3, a pan T-cell marker, showed T-lymphocyte infiltration in regions surrounding HEK-Tom cells as early as 1 week following transplantation (Fig. 7B). This immune reaction to HEK-Tom cells did not require significant tissue damage, since it is present in sham mice that only received a craniotomy (Fig. 7B). CCI resulted in mainly astrogliosis in the corpus callosum without marked changes in the SVZ for GFAP, CDllb, or lymphocyte infiltration (Fig. 7C-E). At 2 weeks post-transplantation, HEK-Tom cells induced a robust, localized reaction in astrocytes (Fig. 7F), macrophages/microglia cells (G), and T-lymphocytes (Fig. 7H). Finally, increased Gli1-YFP labeling was observed in the SVZ adjacent to HEK-Tom cells (Fig. 7I, J). Thus, the robust immune rejection response to human cells activated Shh signaling in endogenous cells in the adjacent regions of the SVZ.

**Figure 7.**
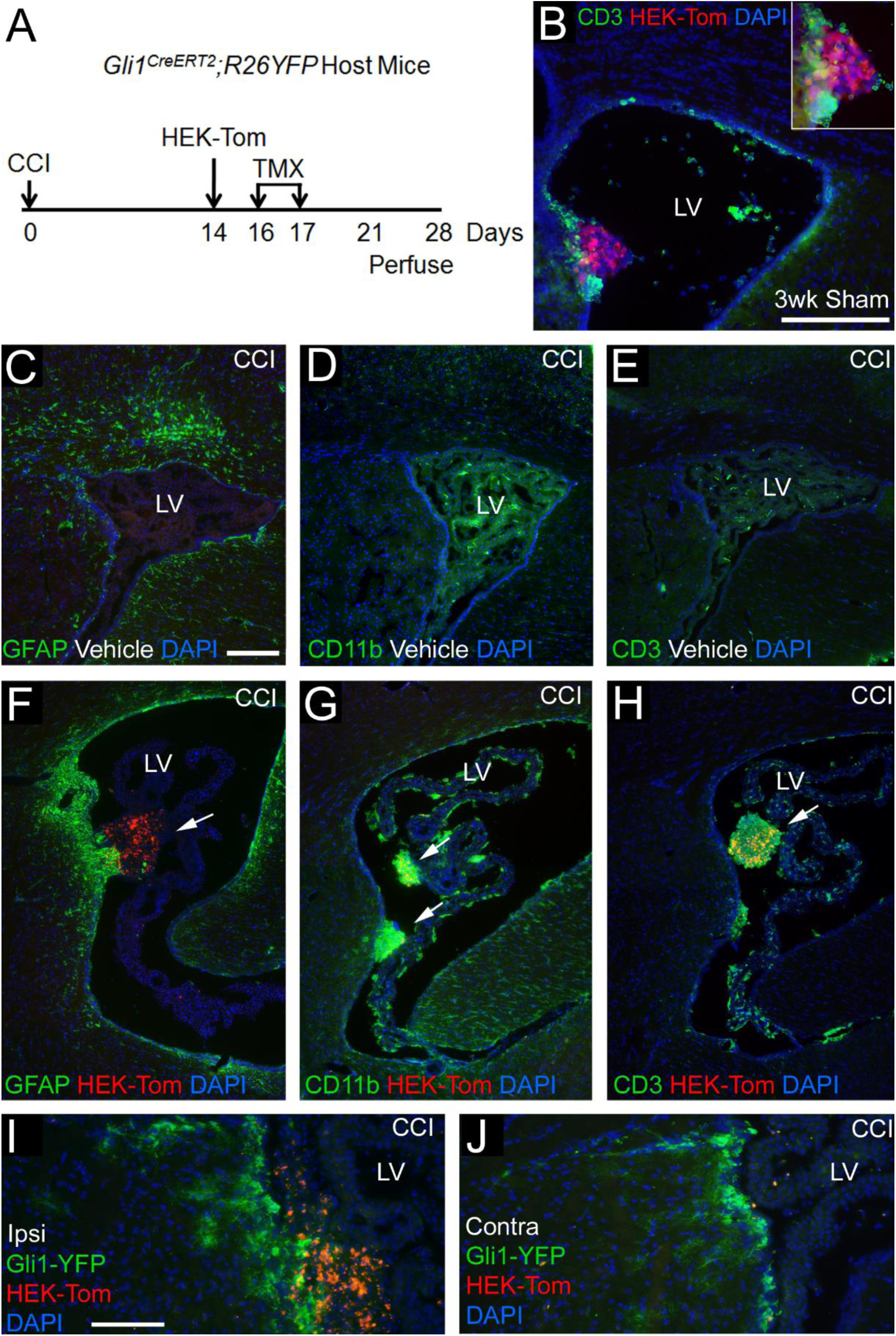
Induced genetic *in vivo* labeling of host cells expressing *Gli1* in the subventricular zone (SVZ) following brain injury and human non-NSC transplantation. **A:** Experimental time line for human HEK transplantation into *Gli1*^*CreERT2*^*;R26YFP* host mice. The HEK293 cells expressing the *tdTomato* reporter (HEK-Tom cells) were transplanted into the lateral ventricle (LV) 2 weeks after craniotomy and the controlled cortical impact (CCI) model of TBI to produce cortical tissue damage. **B:** Immunohistochemistry for the pan T-lymphocyte cell (T-cell) marker CD3 shows T-lymphocytes infiltrate regions surrounding the human HEK293 cells as early as 1 week following transplantation into a sham mouse. **C-H:** Coronal sections from injured mice were immunostained with either GFAP (astrocytes, green), CD11b (macrophages, green), or CD3 (T-cells, green) to analyze the host immune response at 2 weeks post-injection. Immunolabeling for GFAP (C, F), CD11b (D, G) and CD3 (E, H) shows a localized immune response to HEK-Tom cells in the LV (F-H) that is not observed with vehicle injection (C-E). **I-J:** Genetic *in vivo* labeling of host cells with YFP (green) indicates *Gli1* expression after HEK-Tom transplantation. Gli1-YFP labeling is increased in the ipsilateral SVZ adjacent to HEK-Tom cells (I) compared to the contralateral side (J). All sections were counterstained with DAPI (blue) nuclear stain. Scale bar = 200 μm (C, D), 100 μm (I). Representative images are shown from analysis of sham + HEK-Tom (n = 1), CCI + Vehicle (n = 1), CCI + HEK-Tom (n = 2).

## DISCUSSION

NSCs hold therapeutic potential for cell replacement and/or for modulatory interactions with endogenous cells to promote regenerative processes. The current study examined effects of intraventricular transplantation of adult NSCs in experimental TBI. NSCs were transplanted at 2 weeks post-TBI to reflect a clinical time point that could be practical for clinical care decisions and for preparation of immunologically compatible cells for transplantation. Adult NSCs were used as a stem cell source with low risk of tumor formation. A relatively low number of cells was transplanted to minimize mass effects and address the limited amplification capacity that may be likely to occur using protocols to generate autologous NSCs.

Our studies focused on neuroinflammation and neuroregeneration associated with damage in white matter tracts from diffuse axonal injury in TBI. NSCs transplanted into the lateral ventricles survived and remained undifferentiated morphologically as single cells, or small spheres, adhered along the ventricle wall and nestled in the choroid plexus. NSC transplantation had a significant immunomodulatory effect in reducing astrogliosis and microglial/macrophage activation in the corpus callosum after TBI.

Transgenic mouse reporter lines were used to monitor *in vivo* activation of the Shh pathway as a candidate mechanism for transplanted NSC regulation of neuroinflammation as well as neuroregeneration from endogenous stem and progenitor cell populations. Genetic labeling of cells transcribing *Shh* identified neurons as the predominant source of the Shh ligand in the host tissues, which is consistent with prior reports of *Shh* expression [9, 32-34]. Shh labeling was not noticeably altered by TBI or NSC transplantation. TBI did not induce *Shh* expression in reactive astrocytes, activated microglia, or other cells within the corpus callosum.

Using NSCs isolated from transgenic reporter mice, we showed that a subset of transplanted NSCs expressed *Shh in vivo*. Interestingly, *Shh* expression was suppressed in NSCs after TBI, but only among cells in the lateral ventricle region, indicating a specific NSC response based on proximity to the injury. In complementary studies, transgenic reporter mice were used as hosts with genetic labeling of cells transcribing *Gli1*, which is a readout of an active signaling through the canonical Shh pathway. Endogenous cells in the host SVZ exhibited robust *Gli1* labeling. However, NSC transplantation did not activate *Gli1* expression in either the SVZ or in the corpus callosum.

White matter tracts adjacent to the lateral ventricles have been shown to benefit from the effects of intraventricular cell transplantation in experimental models of multiple sclerosis. Neural stem/precursor cell intraventricular transplantation promoted remyelination in experimental autoimmune encephalomyelitis, which produces focal demyelinating lesions in multiple sites throughout the brain and spinal cord [35]. Intraventricular transplantation of syngeneic NSCs reduced astroglial scarring, which may be mediated through down-regulation of factors in the lesion environment that drive reactive astrogliosis, such as fibroblast growth factor [35]. Other studies in this model also showed that neural stem/precursor cells transplanted into the ventricle acted through an anti-inflammatory mechanism to reduce both the innate and adaptive immune response [36]. The use of syngeneic NSCs in these reports, as in the current studies, indicates that immunosuppression is an effect of adult NSC characteristics and is not likely due to factors associated with a general rejection response to the transplanted cells. Using the cuprizone model of corpus callosum demyelination, intraventricular transplantation of NSCs isolated from newborn mice was shown to promote endogenous cell remyelination, which may be mediated through NSC secretion of platelet-derived growth factor and fibroblast growth factor 2 that act as mitogens for oligodendrocyte progenitors [12]. These findings indicate the potential for intraventricular delivery of neural stem/precursor cells to both modulate the immune response and support regenerative mechanisms after white matter injury. Results of the current study are consistent with NSCs suppressing the immune response, but do not provide evidence of transplanted NSC stimulation of a regenerative response in endogenous cells of the host SVZ.

NSCs transplanted into the ventricles may act through direct and indirect mechanisms to modulate expression of regulatory molecules. NSCs can synthesize growth factors *in vivo* after transplantation, as shown for *in vivo* synthesis of Shh in transplanted NSCs in the current experiments with genetic *in vivo* labeling induced from the *Shh* promoter. Transplanted NSCs may also interact with host cells to modulate expression of signaling from endogenous cells of the choroid plexus, ventricle wall, and/or SVZ. However, transplanted NSCs did not appear to induce *Shh* expression in host ependymal or choroid plexus cells in our experiments.

To further explore host-transplant interactions, intraventricular transplantation of human non-NSC cells was used as a positive control to elicit an immune rejection response to then examine the effect on *Gli1* labeling in the host SVZ. T-lymphocytes and macrophages responded strongly to HEK cells transplanted after an invasive form of TBI. Interestingly, this inflammatory response corresponded with localized *Gli1* labeling in adjacent host SVZ cells. Transplanted HEK cells, but not NSCs, also disrupted the normal expression of syndecan-1 in the choroid plexus (data not shown), which indicates HEK cells disrupt the integrity of the blood-cerebrospinal fluid barrier at the choroid plexus that normally prevents immune cell infiltration [37]. Therefore, T-lymphocytes or macrophage activation may be required to induce active Shh signaling. Breakdown of the barriers between the blood and cerebrospinal fluid or brain compartments may also contribute. Consistent with this interpretation, apoptotic/stimulated T-lymphocytes have been reported to release microvesicles containing Shh [38]. In addition, Shh signaling may regulate the integrity of the blood-brain barrier in that inflammation stimulates increased Shh immunolabeling of astrocytes that interact with Gli1 expressing endothelial cells [39]. Overall, our current observations are consistent with previous studies that proposed blood-brain barrier disruption may be required to stimulate Shh signaling in the adult brain, while astrogliosis alone is not sufficient [25, 33]. However, it is not clear yet how Gli1 activation may be influenced by blood-brain barrier disruption. *Gli1* genetic labeling in astrocytes was reduced in lesion areas of the cerebral cortex when TMX was administered after TBI from controlled cortical impact, which results in bleeding from vascular injury [17].

Prior studies from our lab and others have reported a lack of *Gli1* expression in the corpus callosum of naïve adult mice and after TBI [9, 17]. *Gli1* genetic labeling of cells in the corpus callosum was not observed even after microinjection of a Shh pathway agonist into the corpus callosum in naïve adult mice [17]. Indeed, recent studies showed that inhibition of Gli1 may be required for NSCs to be recruited from the SVZ to demyelinated lesions and to subsequently differentiate into oligodendrocytes [40]. This interpretation is also consistent with our prior study showing early transient down regulation of *Gli1 in vivo* labeling in the SVZ in a model of TBI that produces axon damage and demyelination in the corpus callosum [17, 18]. In contrast, studies that used in situ hybridization to detect *Gli1* expression reported an increase of *Gli1* transcripts in corpus callosum lesions after experimental demyelination [13]. The conflicting results could be due to differences in the approaches used to detect Gli1, but also to differences in the inflammatory signals and blood-brain barrier integrity associated with the different pathologies. In addition, although the *Gli1* genetic labeling approach used in the current study is a specific indicator of active *in vivo* signaling through the canonical Shh pathway, Shh can also act through non-canonical mechanisms that do not require Gli1 activation [41, 42].

## CONCLUSIONS

These studies demonstrate that adult NSCs transplanted into the lateral ventricle remain within the lateral and third ventricles for at least two weeks and significantly reduce neuroinflammation in the corpus callosum. These findings are consistent with potential applications for the diffuse white matter injury experienced in TBI. We report that NSC transplantation significantly reduces reactive astrogliosis and microglial/macrophage activation in the corpus callosum after TBI. These studies are also the first to investigate the potential for transplanted NSCs to act through Shh signaling using induced genetic *in vivo* labeling. This genetic approach demonstrated Shh ligand expression *in vivo* in a subset of transplanted NSCs, which was suppressed by TBI. A robust Gli1 response to Shh signaling in endogenous cells of the host SVZ was observed in adult mice. However, NSC transplantation did not stimulate Shh-mediated activation of a regenerative response in the host SVZ. And, NSC reduction of neuroinflammation in the corpus callosum after TBI did not involve Shh signaling through Gli1. Further studies are warranted to better understand the therapeutic potential of transplanted NSCs for suppressing neuroinflammation. Such studies would need to extend to severe forms of TBI and at later time points after injury that will be important for clinical translation and evaluation of the risk-benefit of cell transplantation as a therapeutic intervention.

## Acknowledgements

This work was funded by the Department of Defense in the Center for Neuroscience and Regenerative Medicine (CNRM). We appreciate the technical assistance of Tuan Le and Laurel Beer. We thank Dr. Sohyun Ahn (National Institute for Child Health, Bethesda, MD) for providing the *Gli1*^*CreERT2*^*;R26YFP* and *Gli1*^*CreERT2*^*;R26tdTomato* mice and Dr. Sharon Juliano for providing the C57BL/6-Tg(UBC-GFP)30Scha/J mice. The authors declare that there is no conflict of interest regarding publication of this paper.

## Supplemental Materials

**Figure S1.**
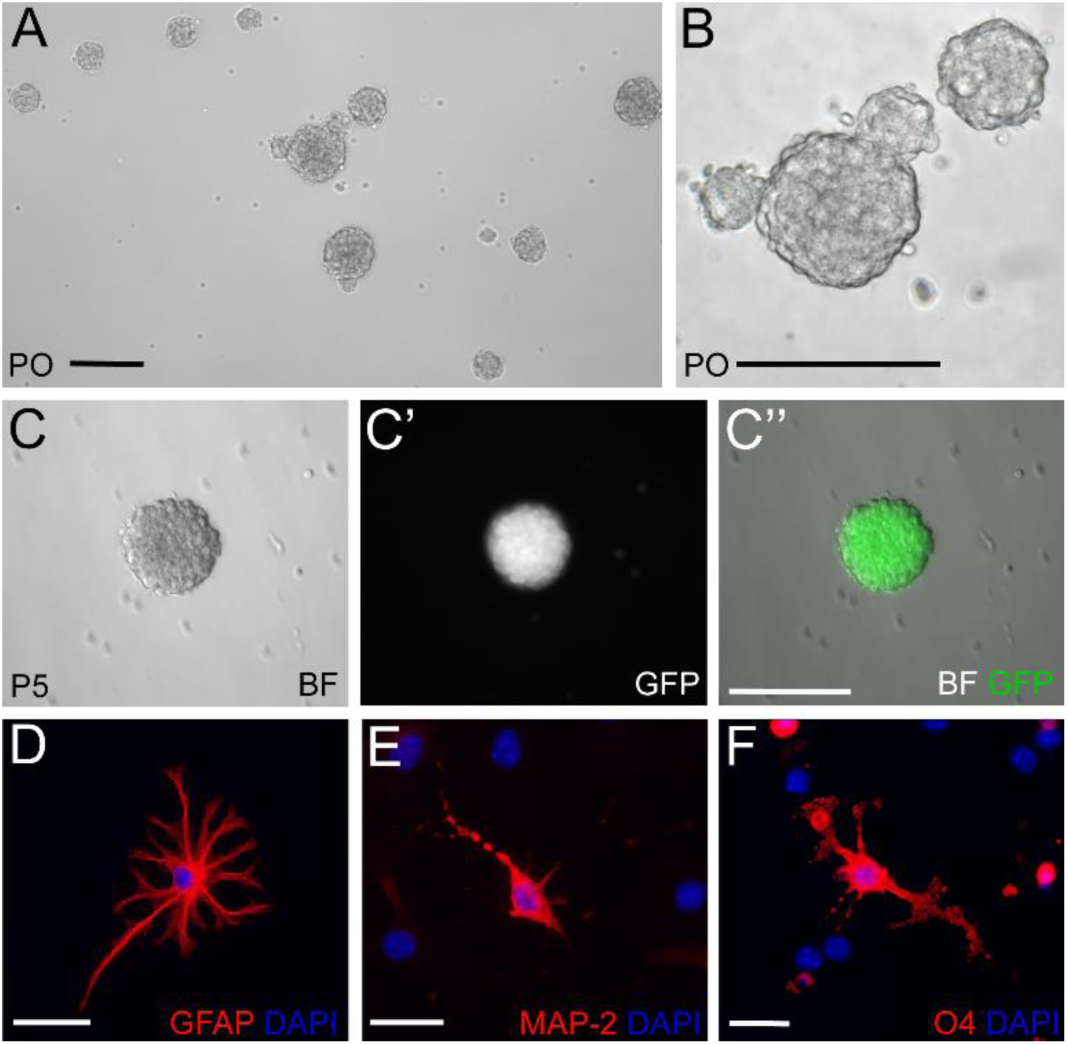
Demonstration of *in vitro* neurosphere formation and multipotentiality of NSCs derived from the subventricular zone (SVZ) of adult mice. **A-C:** Adult NSCs form neurospheres in an *in vitro* suspension culture. Phase contrast image of neurospheres with varying diameters at 3 days after isolation from the SVZ of an adult mouse (A) with higher magnification of middle set (B). NSCs isolated from the SVZ of adult C57BL/6-Tg(UBC-GFP)30Scha/J mice, shown 3 days after passage 5, form neurospheres *in vitro* (C; phase contrast) and express green fluorescent protein (C’; GFP; C”; merge) **D-E:** *In vitro* differentiation shows multipotency of adult NSCs based on immunolabeling for cell type-specific lineage markers of astrocytes (GFAP; D), neurons (MAP-2; E), and oligodendrocytes (O4; F). Scale bars = 100 mm (A, B, C-C”), 40 μm (D), 20 μm (F).

**Table S1.** Mouse lines.

|  | Full name | Source | RRID |
| --- | --- | --- | --- |
| *Shh<sup>CreERT2</sup>* | *B6.129S6-Shh<sup>tm2(cre/ERT2)Cjt</sup>/J* | Jackson Laboratories<br>Cat #005623 | IMSR:JAX_005623 |
| *Gli1<sup>CreERT2</sup>* | *Gli1<sup>tm3(cre/ESR1)Alj</sup>/J* | Jackson Laboratories<br>Cat #007913 | IMSR: JAX_007913 |
| *R26YFP* | *B6.129X1-Gt(ROSA)26Sor<sup>tm1(EYFP)Cos</sup>/J* | Jackson Laboratories<br>Cat #006148 | IMSR:JAX_006148 |
| *R26tdTomato* | *B6.Cg-Gt(ROSA)26<sup>Sortm14(CAG-tdTomato)Hze</sup>/J* | Jackson Laboratories<br>Cat #007908 | IMSR:JAX_007908 |
| *R26mT/mG* | *B6.129(Cg)-Gt(ROSA)26Sor<sup>tm4(ACTB-tdTomato,-EGFP)Luo</sup>/J* | Jackson Laboratories<br>Cat #007676 | IMSR:Jax_007676 |
|  | *C57BL/6-Tg(UBC-GFP)30Scha/J* | Jackson Laboratories<br>Cat #004353 | IMSR: Jax_004353 |

**Figure S2.**
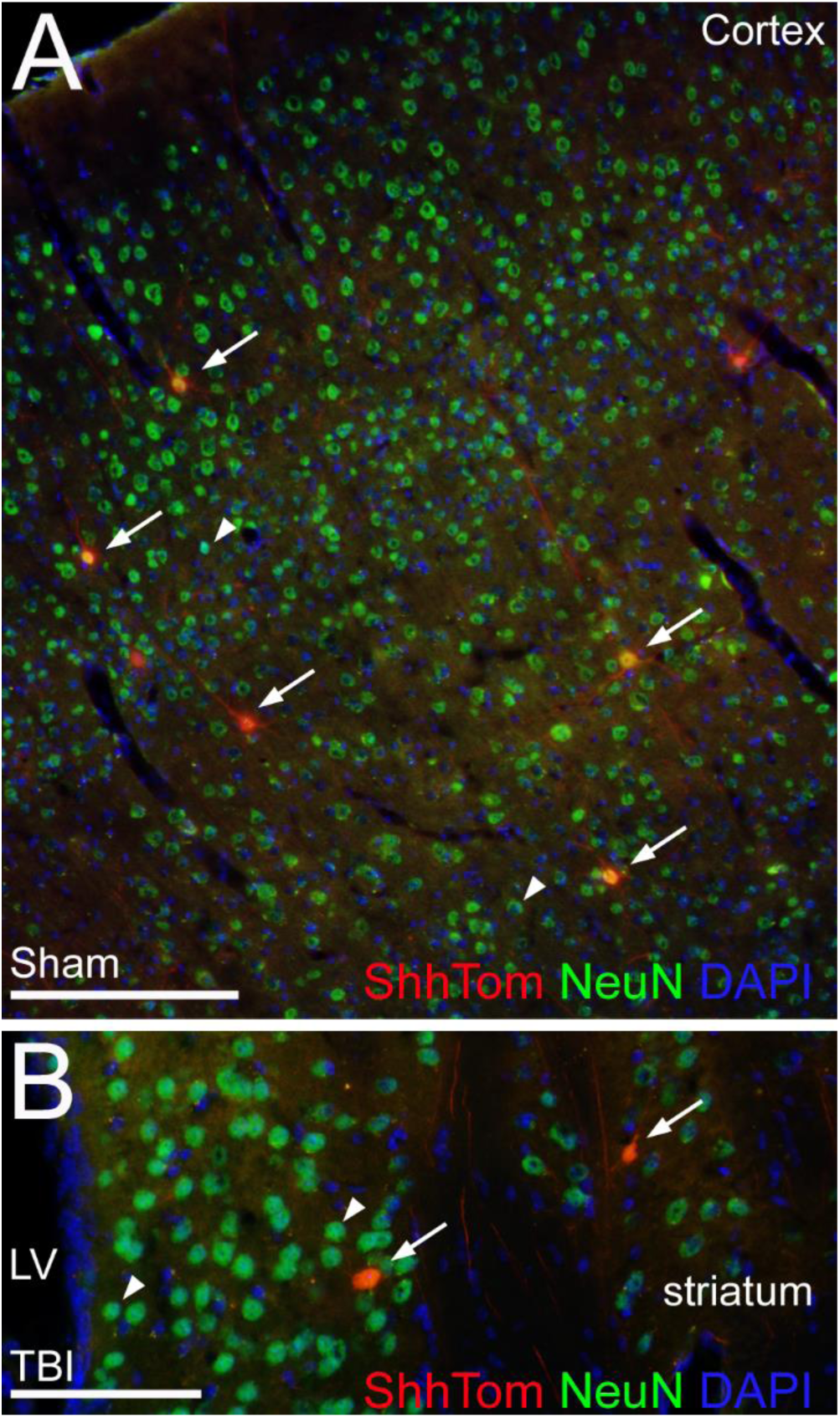
Immunohistochemical identification of Shh-Tom cells as neurons. Coronal sections showing the cerebral cortex (A) and striatum (B) regions from *ShhCreERT2;R26tdTomato* mice. Neurons were identified by expression of the neuronal cell type-specific marker NeuN. Shh-Tom labeled cells (red) exhibit co-labeling with NeuN (green), with the overlap appearing yellow/orange (arrows). Shh-Tom neurons are distributed among cortical layers and are found near the lateral ventricle (LV). Scale bars 200 μm (A), 100 μm (B).

**Figure S3.**
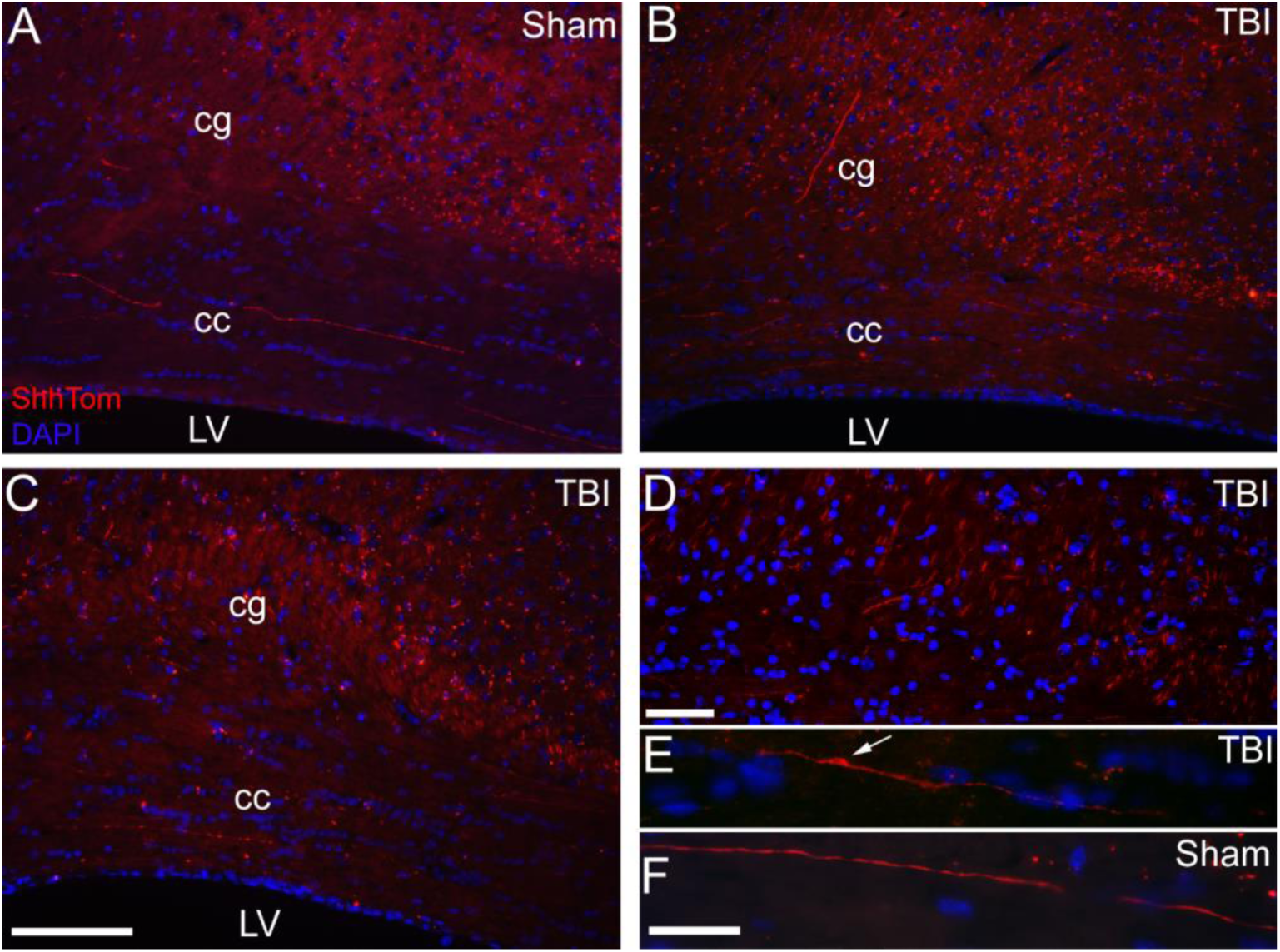
Induced genetic *in vivo* labeling of neurons synthesizing *Shh* that project axons in the corpus callosum and cingulum. Tissues were prepared as in Figure 3, which shows higher magnification examples of transplanted NSC-GFP. **A-C:** Coronal sections showing Shh-Tom labeling (red) with DAPI nuclear stain (blue) in the corpus callosum (cc) and cingulum (cg), regions, which exhibit axon damage and neuroinflammation in this TBI model. The pattern of heritable Shh-Tom labeling of axons in the cingulum and corpus callosum was similar between sham mice (A), TBI mice (B), and TBI mice that received NSC-GFP cells transplanted (B) into the lateral ventricle (LV). **D-F:** Higher magnification of Shh-Tom labeled axons in the cingulum (D) and corpus callosum (E-F). Shh-Tom axons can exhibit greater variation in diameter after TBI (E, arrow) than Shh-Tom axons in sham mice (F). Scale bar = 100 μm (A), 50 μm (D), 25 μm (F). Representative images are shown from analysis of a cohort of sham + NSC-GFP (n = 2), TBI + vehicle (n = 3), and TBI+ NSC-GFP (n = 3) mice.

**Figure S4.**
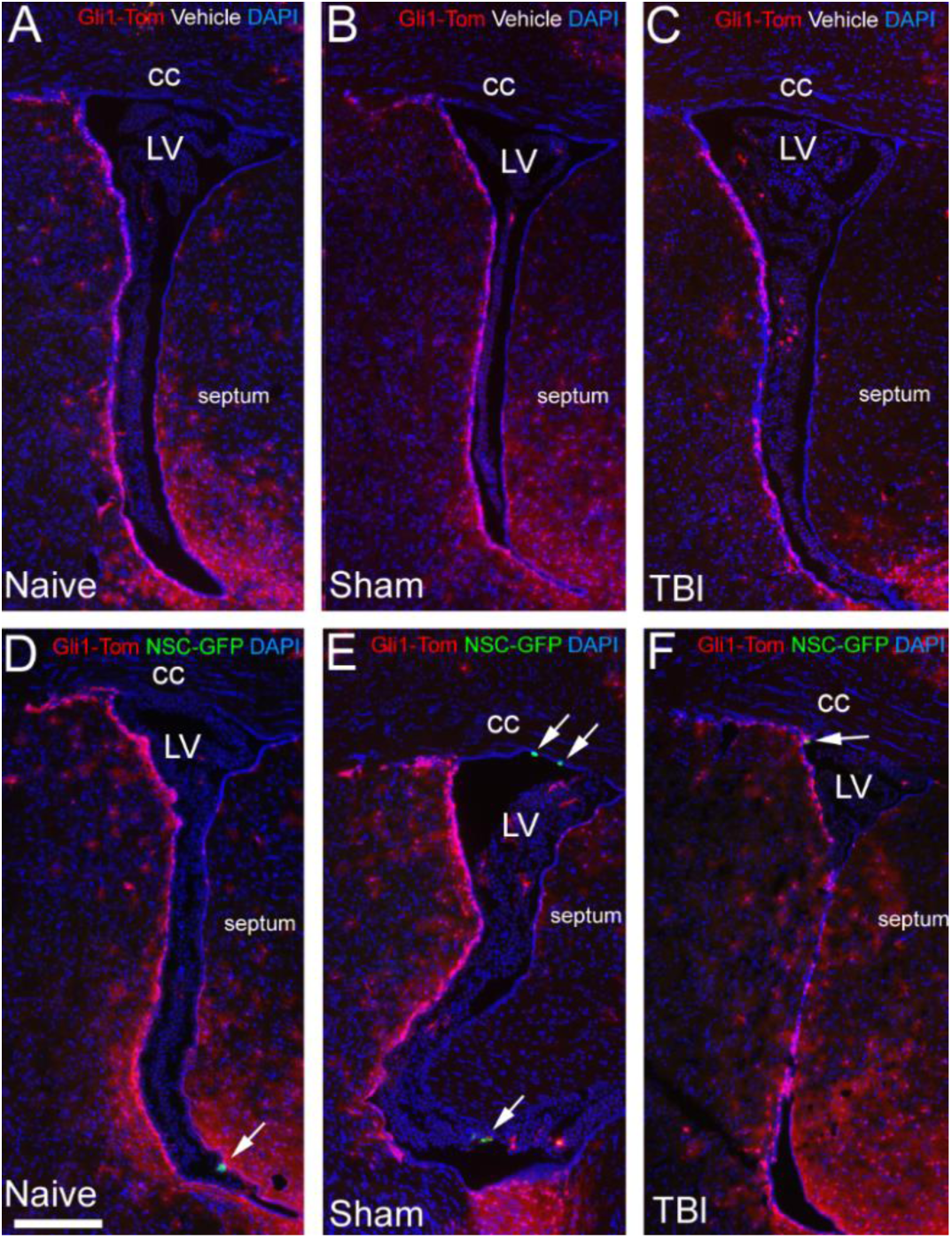
Distribution of Shh responsive cells in regions adjacent to the lateral ventricles. Tissues were prepared as in Figure 5. Coronal sections show Gli1-Tom labeling in endogenous cells of the subventricular zone (SVZ) and regions adjacent to the lateral ventricles (LV). **A-C:** *Glil1CreERT*^*2*^*;R26tdTomato* mice with vehicle injections show Gli1-Tom genetic labeling of cells in the SVZ, septum, and striatum. **D-F:** *Gli1CreERT*^*2*^*;R26tdTomato* mice with transplanted NSC-GFP cells (arrows), identified by constitutive green fluorescent protein, have similar Gli1-Tom expression to vehicle controls. Gli1-Tom labeled cells are not evident in the corpus callosum (cc) regardless of injury or NSC transplantation conditions. Representative images from cohorts of naïve + vehicle (n = 3), naïve + NSC-GFP (n = 2), sham + vehicle (n = 4), sham + NSC-GFP (n = 4), TBI + vehicle (n = 3), and TBI + NSC-GFP (n = 4).

## References

1. Roozenbeek, B., A.I. Maas, and D.K. Menon, Changing patterns in the epidemiology of traumatic brain injury. Nat Rev Neurol, 2013. 9(4): p. 231–6.

2. Smith, D.H., R. Hicks, and J.T. Povlishock, Therapy development for diffuse axonal injury. J Neurotrauma, 2013. 30(5): p. 307–23.

3. Selassie, A.W., et al., Incidence of long-term disability following traumatic brain injury hospitalization, United States, 2003. J Head Trauma Rehabil, 2008. 23(2): p. 123–31.

4. Hill, C.S., M.P. Coleman, and D.K. Menon, Traumatic Axonal Injury: Mechanisms and Translational Opportunities. Trends Neurosci, 2016. 39(5): p. 311–24.

5. Margulies, S., et al., Combination Therapies for Traumatic Brain Injury: Retrospective Considerations. J Neurotrauma, 2016. 33(1): p. 101–12.

6. Drago, D., et al., The stem cell secretome and its role in brain repair. Biochimie, 2013. 95(12): p. 2271–85.

7. Doetsch, F., et al., Subventricular zone astrocytes are neural stem cells in the adult mammalian brain. Cell, 1999. 97(6): p. 703–16.

8. Palma, V., et al., Sonic hedgehog controls stem cell behavior in the postnatal and adult brain. Development, 2005. 132(2): p. 335–44.

9. Garcia, A.D., et al., Sonic hedgehog regulates discrete populations of astrocytes in the adult mouse forebrain. J Neurosci, 2010. 30(41): p. 13597–608.

10. Petrova, R., A.D. Garcia, and A.L. Joyner, Titration of GLI3 repressor activity by sonic hedgehog signaling is critical for maintaining multiple adult neural stem cell and astrocyte functions. J Neurosci, 2013. 33(44): p. 17490–505.

11. Rafuse, V.F., et al., Neuroprotective properties of cultured neural progenitor cells are associated with the production of sonic hedgehog. Neuroscience, 2005. 131(4): p. 899-916.

12. Einstein, O., et al., Transplanted neural precursors enhance host brain-derived myelin regeneration. J Neurosci, 2009. 29(50): p. 15694–702.

13. Ferent, J., et al., Sonic Hedgehog signaling is a positive oligodendrocyte regulator during demyelination. J Neurosci, 2013. 33(5): p. 1759–72.

14. Donega, M., et al., Systemic injection of neural stem/progenitor cells in mice with chronic EAE. J Vis Exp, 2014(86).

15. Giusto, E., et al., Neuro-immune interactions of neural stem cell transplants: from animal disease models to human trials. Exp Neurol, 2014. 260: p. 19–32.

16. Sullivan, G.M., et al., Oligodendrocyte lineage and subventricular zone response to traumatic axonal injury in the corpus callosum. J Neuropathol Exp Neurol, 2013. 72(12): p. 1106–25.

17. Mierzwa, A.J., et al., Comparison of cortical and white matter traumatic brain injury models reveals differential effects in the subventricular zone and divergent Sonic hedgehog signaling pathways in neuroblasts and oligodendrocyte progenitors. ASN Neuro, 2014. 6(5).

18. Mierzwa, A.J., et al., Components of myelin damage and repair in the progression of white matter pathology after mild traumatic brain injury. J Neuropathol Exp Neurol, 2015. 74(3): p. 218–32.

19. Harfe, B.D., et al., Evidence for an expansion-based temporal Shh gradient in specifying vertebrate digit identities. Cell, 2004. 118(4): p. 517–28.

20. Ahn, S. and A.L. Joyner, Dynamic changes in the response of cells to positive hedgehog signaling during mouse limb patterning. Cell, 2004. 118(4): p. 505–16.

21. Ihrie, R.A., et al., Persistent sonic hedgehog signaling in adult brain determines neural stem cell positional identity. Neuron, 2011. 71(2): p. 250–62.

22. Ahn, S. and A.L. Joyner, In vivo analysis of quiescent adult neural stem cells responding to Sonic hedgehog. Nature, 2005. 437(7060): p. 894–7.

23. Srinivas, S., et al., Cre reporter strains produced by targeted insertion of EYFP and ECFP into the ROSA26 locus. BMC Dev Biol, 2001. 1: p. 4.

24. Madisen, L., et al., A robust and high-throughput Cre reporting and characterization system for the whole mouse brain. Nat Neurosci, 2010. 13(1): p. 133–40.

25. Muzumdar, M.D., et al., A global double-fluorescent Cre reporter mouse. Genesis, 2007. 45(9): p. 593–605.

26. Zhou, Q., et al., Valproic acid inhibits neurosphere formation by adult subventricular cells by a lithium-sensitive mechanism. Neurosci Lett, 2011. 500(3): p. 202–6.

27. Zhou, Q., et al., Histone deacetylase inhibitors SAHA and sodium butyrate block G1-to-S cell cycle progression in neurosphere formation by adult subventricular cells. BMC Neurosci, 2011. 12: p. 50.

28. Armstrong, R.C., et al., Absence of fibroblast growth factor 2 promotes oligodendroglial repopulation of demyelinated white matter. J Neurosci, 2002. 22(19): p. 8574–85.

29. Dobson, N.R., et al., Leukemia/lymphoma-related factor regulates oligodendrocyte lineage cell differentiation in developing white matter. Glia, 2012. 60(9): p. 1378–90.

30. Trapp, B.D., et al., Axonal transection in the lesions of multiple sclerosis. N Engl J Med, 1998. 338(5): p. 278–85.

31. Alvarez-Buylla, A. and R.A. Ihrie, Sonic hedgehog signaling in the postnatal brain. Semin Cell Dev Biol, 2014. 33: p. 105–11.

32. Harwell, C.C., et al., Sonic hedgehog expression in corticofugal projection neurons directs cortical microcircuit formation. Neuron, 2012. 73(6): p. 1116–26.

33. Sirko, S., et al., Reactive glia in the injured brain acquire stem cell properties in response to sonic hedgehog. [corrected]. Cell Stem Cell, 2013. 12(4): p. 426–39.

34. Farmer, W.T., et al., Neurons diversify astrocytes in the adult brain through sonic hedgehog signaling. Science, 2016. 351(6275): p. 849–54.

35. Pluchino, S., et al., Injection of adult neurospheres induces recovery in a chronic model of multiple sclerosis. Nature, 2003. 422(6933): p. 688–94.

36. Pluchino, S., et al., Neurosphere-derived multipotent precursors promote neuroprotection by an immunomodulatory mechanism. Nature, 2005. 436(7048): p. 266–71.

37. Zhang, X., et al., Syndecan-1, a cell surface proteoglycan, negatively regulates initial leukocyte recruitment to the brain across the choroid plexus in murine experimental autoimmune encephalomyelitis. J Immunol, 2013. 191(9): p. 4551–61.

38. Martinez, M.C., et al., Transfer of differentiation signal by membrane microvesicles harboring hedgehog morphogens. Blood, 2006. 108(9): p. 3012–20.

39. Alvarez, J.I., et al., The Hedgehog pathway promotes blood-brain barrier integrity and CNS immune quiescence. Science, 2011. 334(6063): p. 1727–31.

40. Samanta, J., et al., Inhibition of Gli1 mobilizes endogenous neural stem cells for remyelination. Nature, 2015. 526(7573): p. 448–52.

41. Brennan, D., et al., Noncanonical Hedgehog signaling. Vitam Horm, 2012. 88: p. 55–72.

42. Robbins, D.J., D.L. Fei, and N.A. Riobo, The Hedgehog signal transduction network. Sci Signal, 2012. 5(246): p. re6.

